# Chromosome-level genome assembly of the horned-gall aphid, *Schlechtendalia chinensis* (Hemiptera: Aphididae: Erisomatinae)

**DOI:** 10.1101/2021.02.17.431348

**Authors:** Hong-Yuan Wei, Yu-Xian Ye, Hai-Jian Huang, Ming-Shun Chen, Zi-Xiang Yang, Xiao-Ming Chen, Chuan-Xi Zhang

**Author notes:** Correspondence Zi-Xiang Yang, Research Institute of Resource Insects, Chinese Academy of Forestry, Kunming, China., Xiao-Ming Chen, Research Institute of Resource Insects, Chinese Academy of Forestry, Kunming, China., Chuan-Xi Zhang, Institute of Insect Sciences, Zhejiang University, Hangzhou, China. Contributed equally (Hong-Yuan Wei, Yu-Xian Ye). Funding information National Natural Science Foundation of China, Grant/Award number: 31872305, U1402263; The basic research program of Yunnan Province, Grant/Award number: 202001AT070016; The grant for Innovative Team of ‘Insect Molecular Ecology and Evolution’ of Yunnan Province.

## Abstract

The horned gall aphid *Schlechtendalia chinensis*, is an economically important insect that induces galls valuable for medicinal and chemical industries. *S. chinensis* manipulates its host plant to form well-organized horned galls during feeding. So far, more than twenty aphid genomes have been reported; however, all of those are derived from free-living aphids. Here we generated a high-quality genome assembly of *S. chinensis*, representing the first genome sequence of a galling aphid. The final genome assembly was 280.43 Mb, with 97% of the assembled sequences anchored into thirteen chromosomes. *S. chinensis* presents the smallest aphid genome size among available aphid genomes to date. The contig and scaffold N50 values were 3.39 Mb and 20.58 Mb, respectively. The assembly included 96.4% of conserved arthropod and 97.8% of conserved Hemiptera single-copy orthologous genes based on BUSCO analysis. A total of 13,437 protein-coding genes were predicted. Phylogenomic analysis showed that *S. chinensis* formed a single clade between the *Eriosoma lanigerum* clade and the Aphidini+Macrosiphini aphid clades. In addition, salivary proteins were found to be differentially expressed when *S. chinensis* underwent host alternation, indicating their potential roles in gall formation and plant defense suppression. A total of 36 cytochrome P450 genes were identified in *S. chinensis*, considerably fewer compared to other aphids, probably due to its small host plant range. The high-quality *S. chinensis* genome assembly and annotation provide an essential genetic background for future studies to reveal the mechanism of gall formation and to explore the interaction between aphids and their host plants.

## 1 Introduction

Many aphid species are economically important plant pests that feed on plant sap and also caused damage by transmitting plant viruses. Around 100 aphid species have been identified as significant agricultural pests among the approximately 5,000 known species (Blackman & Eastop, 2017). Up to this point, studies on aphid genomes have mostly focused on the subfamily Aphidinae (International Aphid Genomics Consortium, 2010; Li et al., 2019; Mathers, 2020; Mathers et al., 2017; Mathers, Mugford, et al., 2020; Mathers, Wouters, et al., 2020; Nicholson et al., 2015; Thorpe et al., 2018; Wenger et al., 2016). Genome sequencing efforts of other subfamilies, however, such as Eriosomatinae that are distantly related to Aphidinae are very limited (Julca et. al. 2020; Biallo et al., 2020). Unlike most free-living aphids, the galling species from Eriosomatinae induce the formation of galls on their primary host plant and live in the interior. Galling aphids may serve as good models for the study of unique ecological and behavioral phenomena for insect-plant interaction and coevolution (Moran, 1989; David Wool, 2004).

The horned gall aphid, *Schlechtendalia chinensis* (Hemiptera: Aphididae: Eriosomatinae: Fordini), is one of the most economically valuable insects, since it induces the formation of gallnuts that have been used for medicinal purposes and in chemical industries. The components of gallnuts are useful or essential gradients in producing inks, wine, food, cosmetic antioxidants, and animal feed. These widely uses are due to their high tannin levels which range from 50% to 70% (Zhang, Tang, & Cheng, 2008). The annual yield of gallnuts in China is 8,000-10,000 tons, accounting for >90% of the total yield worldwide (Zhang, Tang, & Cheng, 2008). *S. chinensis* shows a peculiar complex life cycle involving both sexual and asexual reproduction stages with a host alternation between the Chinese sumac (*Rhus chinensis*, Anacardiaceae) and mosses (*Plagiomnium spp*. Mniaceae). In this holocyclic life cycle, a fundatrix produced by a mating female crawls along the trunk and feeds on a new leaf, where it induces the formation of a horned-gall. The fundatrix then continues to feed inside the gall and reproduces fundatrigeniae via parthenogenesis. From here on, the gall grows quickly and forms a highly organized structure. During gall development, drastic morphological rearrangements occur in leaf tissues. For example, the palisade tissues of a gall are reorganized and replaced by parenchym cells (Liu *et al*., 2014). In autumn, galls become mature and burst to open, alate autumn migrants fly to nearby mosses and reproduce nymphs for overwintering. In the following spring, nymphs develop in the moss into spring migrants that fly back to the primary host, *chinensis*. Here they produce female and male offspring (sexuales) with degenerated mouthparts that allow them to attach themselves to the trunk crevices. After mating, each female reproduces only a fundatrix, starting the next life cycle all over again (Figure 1) (Zhang, Qiao, Zhong & Zhang, 1999; Blackman and Eastop, 2000). Taken together, *S. chinensis* participates in a series of all-female parthenogenetic generations and only a single sex generation. This unusually life cycle with various morphologically distant aphids at different stages is likely driven by adaptation to different environmental conditions. Unlike most free-living aphids from the Aphidinae taxon, galling aphids have many distinct biological characteristics. The most striking characteristic is that the feeding of most galling aphids does not seriously affect the health of their host plants. In fact, the galls of *S. chinensis* even provide some benefits to its primary host plant (Chen et al., 2020).

**Figure 1.**
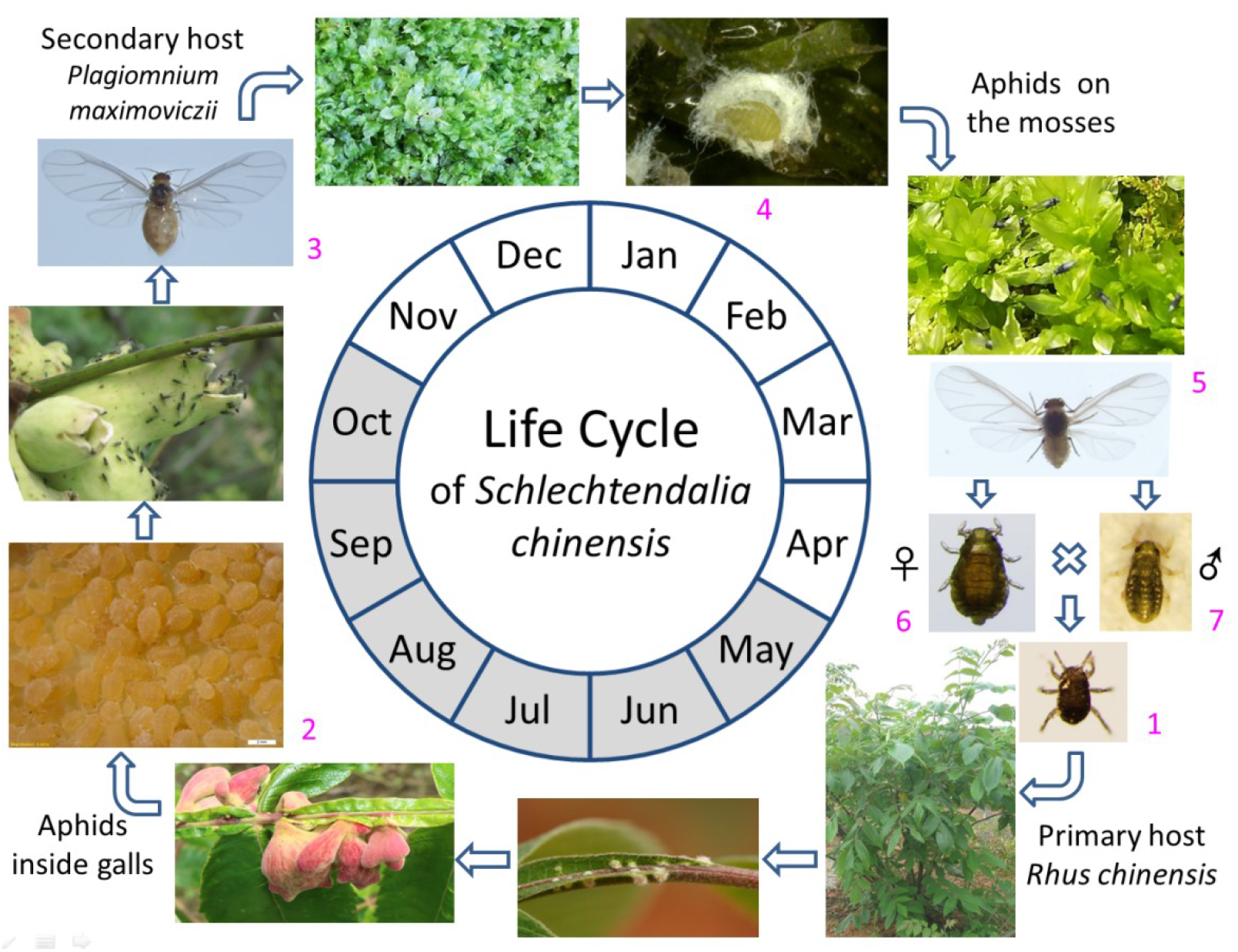
Life cycle diagram of *Schlechtendalia chinensis*. A typical life cycle of the horned gall aphid in Hubei, China. A fundatrix (1) find a tender leaf on the primary host *Rhus chinensis*, to feed and initialize gall formation, and feeds inside the induced gall by the end of April or the beginning of May. Afterwards, the fundatrix and her wingless daughters (called fundatrigeniae) (2) reproduce for generations viviparously and parthenogenetically within the gall from May to October. The gall size increases gradually along with the growth of the aphid population in it. At the end of October, winged autumn migrants (3) emerge from the gall and fly away after the gall opened. The migrants find the moss *Plagiomnium maximoviczii* nearby where they produce nymphal offspring (4). The nymphs feed on the moss and secrete waxes to wrap themselves up for overwintering. Winged spring migrants (5) emerge by the end of March, then fly back to the primary host and reproduce sexual females (6) and males (7) in the trunk crevices. After mating, the female reproduce a fundatrix to begin the next life cycle. *Graphs not in scale. Stippled sector indicating the in-gall stages.

The complexity both in its developmental process and in the structure of its induced galls implies that *S. chinensis* may have unique gene sets that regulate its development and manipulate its host plants (Takeda et al., 2019; Hirano et al., 2020). The underlying molecular mechanisms for its complex life cycle and ability to parasitize on host plants via apparently harmless galls remain largely unknown. Here we generated a high-quality chromosome-level genome assembly of *S. chinensis*, representing the first genome sequence of aphids that induces the formation of closed galls, using a combination of Illumina, Pacific Bioscience (PacBio) and Hi-C technologies. Subsequent, gene prediction, functional annotation and phylogenetic analysis were also carried out to determine the relationship between *S. chinensis* and other aphid species. This reference genome sequence provides information about genomic organization in the Eriosomatinae subfamily and allows comparative genomic studies for a better understanding of genes associated with its own developmental regulation as well as its ability to induce gall formation, living on alternative host plant species and suppress plant defence during parasitism.

## 2 Materials and Methods

### 2.1 Sample collection

*chinensis* samples were collected in October, 2019, from fresh mature galls on *R. chinensis*, in Wufeng county (30°10′ N,110°52′ E,960 m above sea level), Hubei Province, China. Winged and wingless fundatrigeniae in a gall were treated as the same clone. Fundatrigeniae contained within a gall were poured onto a petri dish after dissecting the gall. Impurities such as waxes were removed manual. The fundatrigeniae were then transferred to Eppendorf tubes. Samples were separated into groups with each group containing about 20 individuals. The samples were immediately frozen in liquid nitrogen for 2 hours and then stored at −80°C for later genome, RNA-seq and Hi-C analyses respectively.

In addition to the original samples, a colony was established through artificial cultivation for other genetic studies. To establish a colony, autumn migrants of *S. chinensis* were collected from mature galls and transferred to a nursery of the moss *Plagiomnium maximoviczii*, which were maintained in a greenhouse. In the following year, spring migrants (sexuparae) were collected from the mosses and cultivated in lab. Aphids were transferred to host trees after fundatrix emergence for gall induction. Aphid samples of different stages including fundatrix, fundatrigeniae, autumn migrants, overwinter nymphs, and spring migrants as well as male and female sexuales were collected separately. All aphid samples were immediately frozen in liquid nitrogen for two hours and then stored at −80°C until further analysis.

### 2.2 Genomic sequencing

Genomic DNA was isolated from aphid samples using a DNeasy Blood & Tissue Extraction Kit (Qiagen Inc., Valencia, CA, USA) by following the manufacturer’s instructions. After the measurement of quality and quantity, the DNA samples were used for making a paired-end sequencing library (150 bp in length). The library was sequenced using the Illumina NovaSeq6000 plantform. In addition, a 20 kb library was constructed and sequenced using the PacBio RSII platform at Annoroad Gene Technology Co., Ltd. (Beijing, China).

Hi-C libraries were also constructed from aphid samples according to the Proximo Hi-C procedure and its quality and concentration were determined via an Agilent 2100 Bioanalyzer and qPCR. Briefly, aphid samples were mixed with 1% formaldehyde for 10 min at room temperature and the nuclei were extracted and permeabilized. After the quality of the library was ascertained, different libraries were pooled to achieve required concentrations for Illumina sequencing (Rao et al., 2014).

RNA samples were extracted from the fundatrix, the fundatrigeniae, autumn migrants, nymphs, spring migrants (sexuparae), male and female sexuales, respectively, following the standard protocols provided by the manufacturer. RNA quantity, purity and integrity were determined by a NanoPhotometer and the Agilent 2100 Bioanalyzer. cDNA libraries were built following the chain specific method. The libraries were initially quantified with qubit 2.0 fluorometer and diluted to 1.5 ng/ul. Different libraries were pooled according to the requirements of effective concentration and target data volume for Illumina sequencing.

### 2.3 Genome assembly

We selected a k-mer length of 19 bases and used Illumina paired end reads for k-mer analysis to estimate the genome size and generate k-mer distribution as described by Jellyfish and GenomeScope (Ranallo-Benavidez, Jaron, & Schatz, 2020). We used PacBio sequencing to assembly genome using Wtdbg2. Long reads were used to fix base errors in sequences generated by noisy long reads using NextPolish (Hu, Fang, Su, & Liu, 2019). Pilon was used to improve the draft assembly from Illumina reads (Walker et al., 2014). Finally, HaploMerger2 was used to remove the heterozygosity (Huang, Kang, & Xu, 2017).

Haploid contigs were scaffolded using the 3D *de novo* assembly (3D-DNA) software (Dudchenko et al., 2017). Hi-C sequences were aligned with the draft genome assembly using Juicer. An initial assembly was generated via a 3D-DNA analysis. The initial assembly was validated using JBAT Juicebox, resulting in a finally chromosome-level genome assembly.

### 2.4 Chromosome staining

Salivary glands were dissected from fundatrigeniae of *S. chinensis* in a phosphate-buffer saline (PBS) solution (137 mM NaCl, 2.68 mM KCl, 8.1 mM Na_2_HPO_4_ and 1.47 mM KH_2_PO_4_ at pH 7.4) under a stereomicroscope (COIC, Chongqing, China). The samples were fixed in 4% paraformaldehyde (Sangon Biotech, Shanghai, China) for 1 h and washed with PBS three times. Then the samples were triturated mechanically, and 20 ml DAPI solution (Abcam, Cambridge, USA) was added to stain the chromosomes. Fluorescence images were examined and photographed under a Leica confocal laser-scanning microscope (Leica, Heidelberg, Germany).

### 2.5 Gene annotation

Repetitive sequences were identified based on homology and *de novo* approaches. For the homology-based approach, RepeatMasker was used to screen the *S. chinensis* genome against the Hemiptera ch repeat database, set this parameter to RepeatMasker -pa 4 -e ncbi -species Hemiptera ch -dir. Repeatmodeler identifies repetitive sequences based on *de novo* approaches prediction (Flynn et al., 2020). Statistical results of RepeatMasker and Repeatmodeler analyses were combined.

Gene structure predictions were performed using homology-based, *de novo*, and RNA-seq assisted methods. For the homology-based method, gene structures were predicted based on homology to those from *Eriosoma lanigerum, Acyrthosiphon pisum, Myzus persicae, Aphis glycines, Rhopalosiphum maidis* (Birney, Michele, & Durbin, 2004). For *de novo* prediction, Augustus was first used for Augustus training, then for RNA-seq data to make hint files, and finally run to perform gene prediction (Stanke et al., 2006; Blanco, Parra & Guigó, 2007). In addition, GeneMark-ET, BRAKER, Maker and SNAP were also applied separately to generate ab initio gene predictions. For the RNA-seq assisted method, RNA-seq data were alignedto the *S. chinensis* genome using PASA. Finally, gene prediction results were integrated by EVidenceModeler (Haas et al., 2008).

We aligned the genes to seven functional databases to annotate genes in the *S. chinensis* genome. The databases used in the study were NCBI Non-Redundant Protein Sequence (Nr), Non-Redundant Nucleotide Sequence Database (Nt), SwissProt, Cluster of Orthologous Groups for eukaryotic complete genomes (KOG), The Integrated Resource of Protein Domains and Functional Sites (InterPro), Conserved Domain Database (CDD), Gene Ontology (GO), Kyoto Encyclopedia of Genes and Genomes, Orthology database (KEGG) and evolutionary genealogy of genes: Non-supervised Orthologous Groups (eggNOG). A localized Blast2GO database was built first and then GO annotation was processed via Blast2GO. KAAS of KEGG was used to annotate *S. chinensis* genome sequence quickly, and the pattern of BBH was chosen.

### 2.6 Non-coding RNA identification

Transfer RNAs (tRNAs) were identified using the tRNAscan-SE program (set to default parameters, the same as below unless specific notes). RNAmmer was used to identify 5S/ 8S, 16S/ 18S and 23S/ 28S rRNA. Ribosomal RNAs (rRNAs), microRNAs (miRNAs) and small nuclear RNAs (snRNAs) were identified based on the Rfam database (12.2) (Kalvari et al., 2017).

### 2.7 Phylogenetic analysis

We constructed phylogenetic trees using whole-genome sequences of *S. chinensis* and whole-genome sequences of other ten aphid species including *Cinara cedri, E. lanigerum, Ap. glycines, Ap. gossypii, R. maidis, R. padi, Ac. pisum, Diuraphis noxia, M. cerasi* and *M. persicae*. The aphid genome sequence and gene structure annotation files were downloaded from the NCBI genome database, and genes containing mRNA information were retained and the tail CDS was modified. Finally, the protein and CDS sequences of all genes were obtained. The obtained protein sequences were filtered and analyzed by all vs all blast. Homologous genes were clustered using OrthoMCl (Li, Stoeckert & Roos, 2003). The coding sequences of homologous single copy genes were extracted. Multi-sequence alignments of homologous single copy genes were performed using MAFFT (Katoh, Misawa, Kuma, & Miyata, 2002; Katoh & Standley, 2013). Protein sequence alignments were converted to codon sequence alignments and conserved blocks were extracted from the alignments using Gblocks. The extracted codon sequence alignments of all homologous single copy genes were integrated for phylogenetic trees using RAxML (Castresana, 2000). The branch length of homologous genes was analyzed using PAML, and then compared with the standard tree to eliminate abnormal genes. Then the tree was built using RAxML again (Stamatakis, 2014). By providing the root number and multiple sequence alignment results with calibration point information, the species divergence time can be calculated using mcmctree (a part of the PAML software), and the divergence time within the evolutionary tree can be obtained with 95% confidence (Yang, 2007). The nodal dates of *Ac. pisum* and *M. cerasi* were 42.4-48.0 MYA, and those of *Ac. pisum* and *Ap. gossypii* were 34-60 MYA.

### 2.8 Gene family expansion and contraction

We used CAFÉ to analyze the gene family expansion and contraction by comparing our data with those from 10 other aphids (*C. cedri, E. lanigerum, Ap. glycines, Ap. gossypii, R. maidis, R. padi, Ac. pisum, D. noxia, M. cerasi* and *M. persicae*). Briefly, the quantitative information of gene families of 12 aphids was obtained according to OrthoMCL results. The number of gene families in each species and the trees with divergence time were used as the input information of CAFÉ. Then the expansion and contraction of gene families were measured by CAFÉ (Hahn, Demuth & Han, 2007).

### 2.9 Identification of genes potentially involved in gall formation and host manipulation

A total of 141 proteins have been identified from saliva and dissected salivary glands of *S. chinensis* in a previous study (Yang et al., 2018). Genes coding these 141 proteins were presented in the newly annotated *S. chinensis* genome of this study. The expression of the 141 genes was analyzed based on RNA-seq data. Putative P450 genes from the 10 aphid species were counted and the rates to each total genome were calculated, respectively. An HMM file of the P450 gene family was downloaded and analyzed for Pfam (http://pfam.xfam.org/). The Pfam domain of P450s is PF00067. Then HMMER was used to blast the HMM file of detoxification genes with insect protein sequences in order to find candidate sequences of detoxification gene families in insect protein sequences (score>50, E-value>1E-5). Phylogenetic analysis of P450 genes in *Ac. pisum* and *S. chinensis* were conducted by aligning proteins using ClustalW and then MEGA (v7.0) based on the maximum likelihood method.

## 3 Results and Discussion

### 3.1 Genome sequencing and *de novo* assembly

Sequencing of the fundatrigenia genome using Illumina paired-end technology generated 73.85 Gb raw reads. The k-mer (K=19) analysis indicated that the heterozygosity of *S. chinensis* was 0.89% and the estimated size was 280.43 Mb. This represents the smallest aphid genome among the most recent available aphid genomes (Table S2, S4; Figure S1) (Wenger et al., 2020; Quan et al., 2019). In addition, sequencing of the fundatrigenia genome using the PacBio PS II platform generated 130 Gb of raw data with an N50 read length of 21.03 kb. After quality control, 127,473,721,171 bp were identified from 9,286,929 reads and an N50 read length was 20.58 kb. The raw contig-level assembly comprised 304,774,269 bp with 1,409 contigs, and the contig N50 was 2,961,835 bp, which showed high-quality of this genomic assembly (Table 1).

**Table 1.**
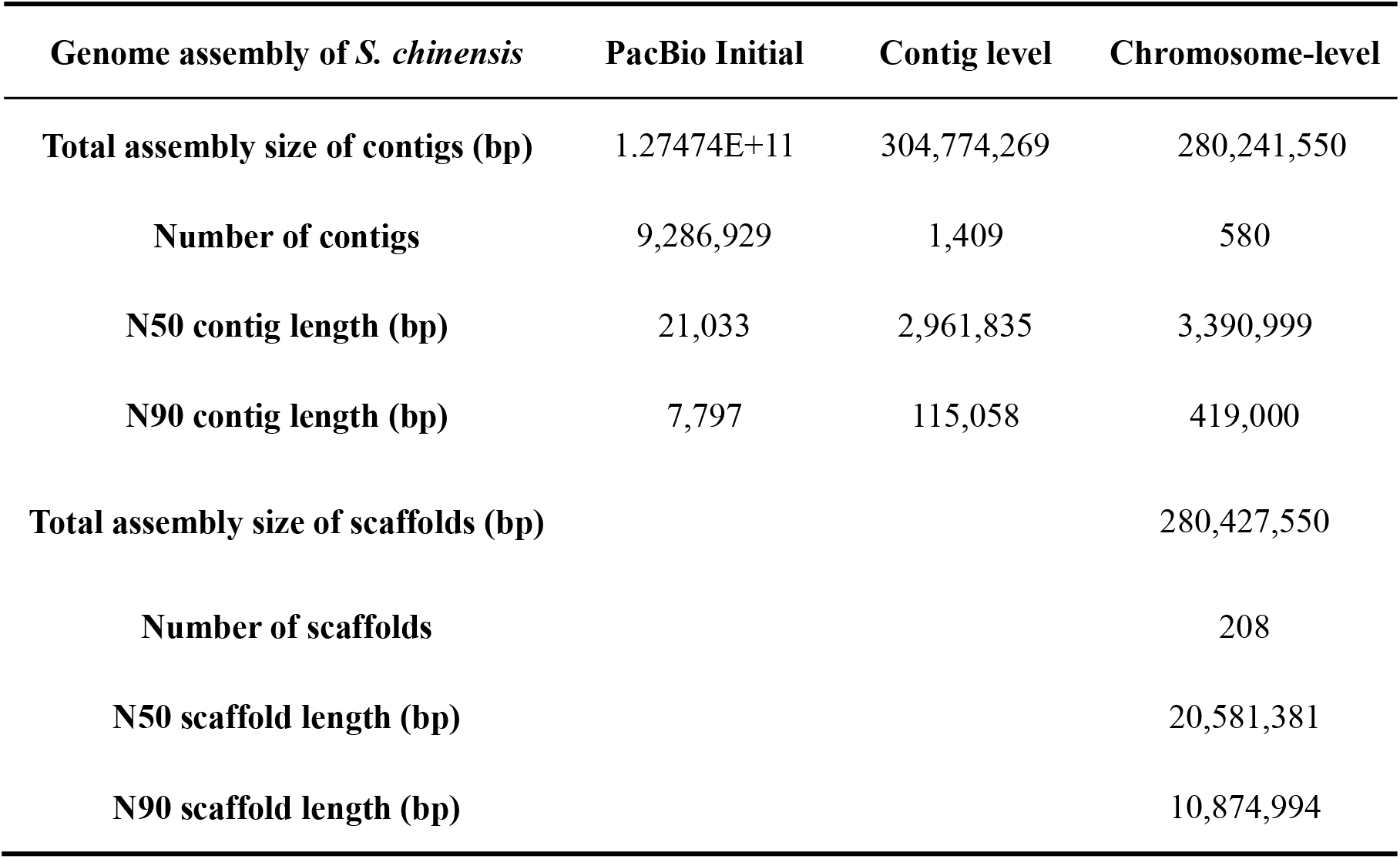
Statistics of the *S. chinensis* genome assembly.

Then, Hi-C data was used to improve our genome assembly. A total of 100.09 Gb (96.23 Gb clean data) (Table S1). The chromosome-level genome was assembled into a total length of 280,427,550 bp and 208 scaffolds using PacBio and Hi-C data. Its contig and scaffold N50 reached 3.39 Mb and 20.58 Mb, respectively (Table 1, S4) (Mathers et. al. 2020). More than 97% of the total genome bases were successfully anchored to 13 chromosomes, with the chromosomic lengths ranging from 5.67 Mb to 86.7 Mb (Table S5; Figure 2). Chromosomic staining was performed to validate the Hi-C analysis result, and 13 chromosomes were identified in the fundatrigenia of *S. chinensis* (Figure S2).

**Figure 2.**
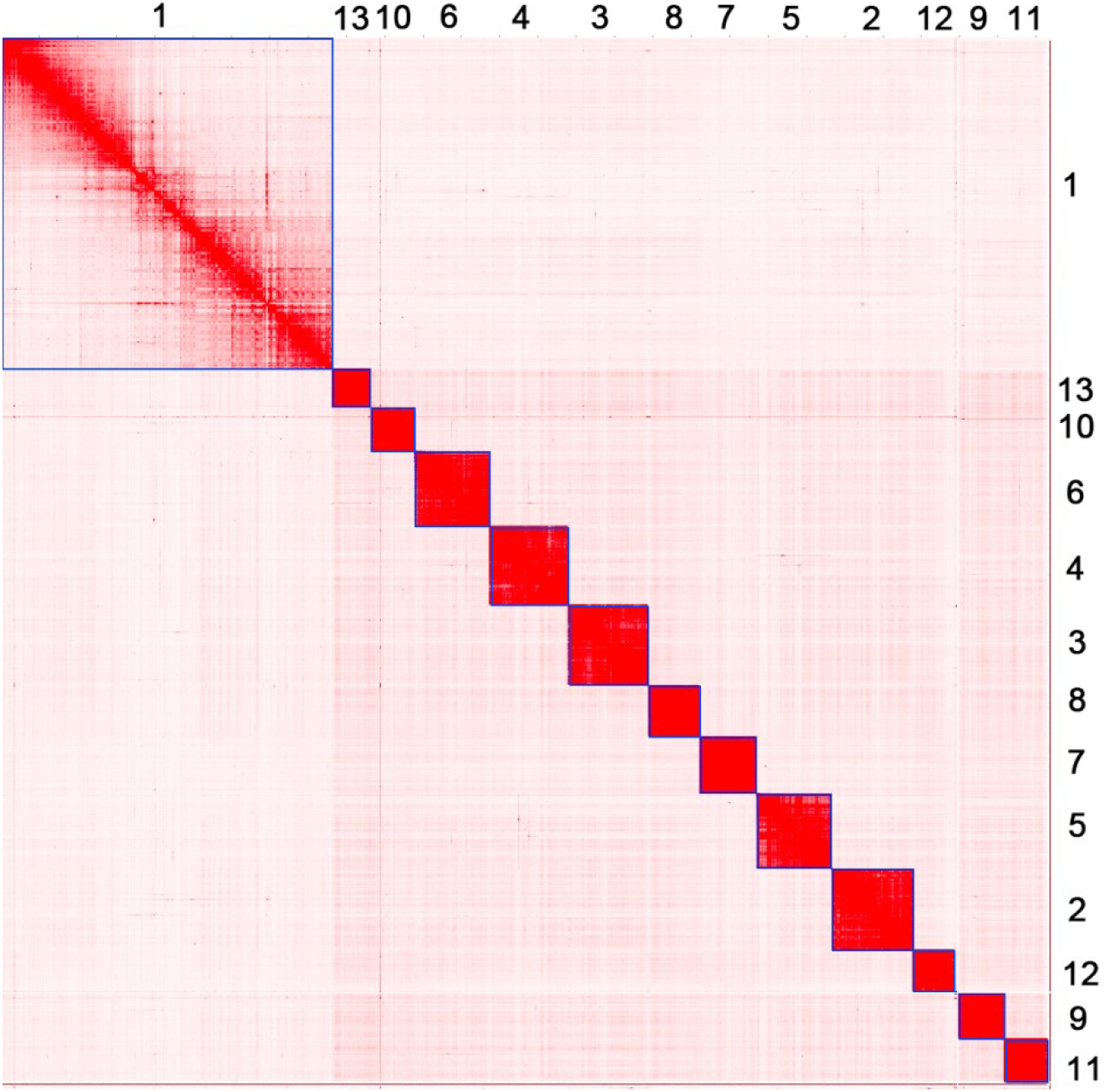
Contact maps of Hi-C interactions among chromosomes in the *S. chinensis* genome. The heatmap was generated by Juicebox software using *in situ* Hi-C data.

In order to evaluate the completeness of the genome assembly, BUSCO analyses against Eukaryota, Arthropoda, Insecta and Hemiptera datasets were performed. The results showed that 91.8%, 96.4%, 94.6% and 97.8% of BUSCO genes could be identified in the *S. chinensis* genome in comparison with the Eukaryota, Arthropod, Insecta and Hemiptera datasets, suggesting the completeness and high quality of our genome assembly (Table S3).

### 3.2 Genome annotation

A total of 162,500,122 bp repetitive sequences were obtained in the *S. chinensis* genome and the proportion of repeats was 57.94% (Table 3 and Figure S6). In total, the number of predicted protein-coding genes was 13,437, which is similar to what has been reported for most aphid species but represents a far lower gene number compared to the genomes of *M. cerasi, E. lanigerum*, and *R. padi*. The average CDS length, exons number per gene, exon length and intron length were 1,853.73 bp, 7.24, 256.19 bp and 822.56 bp, respectively, which are comparable to the corresponding metrics from other aphids (Table 2).

**Table 2.**
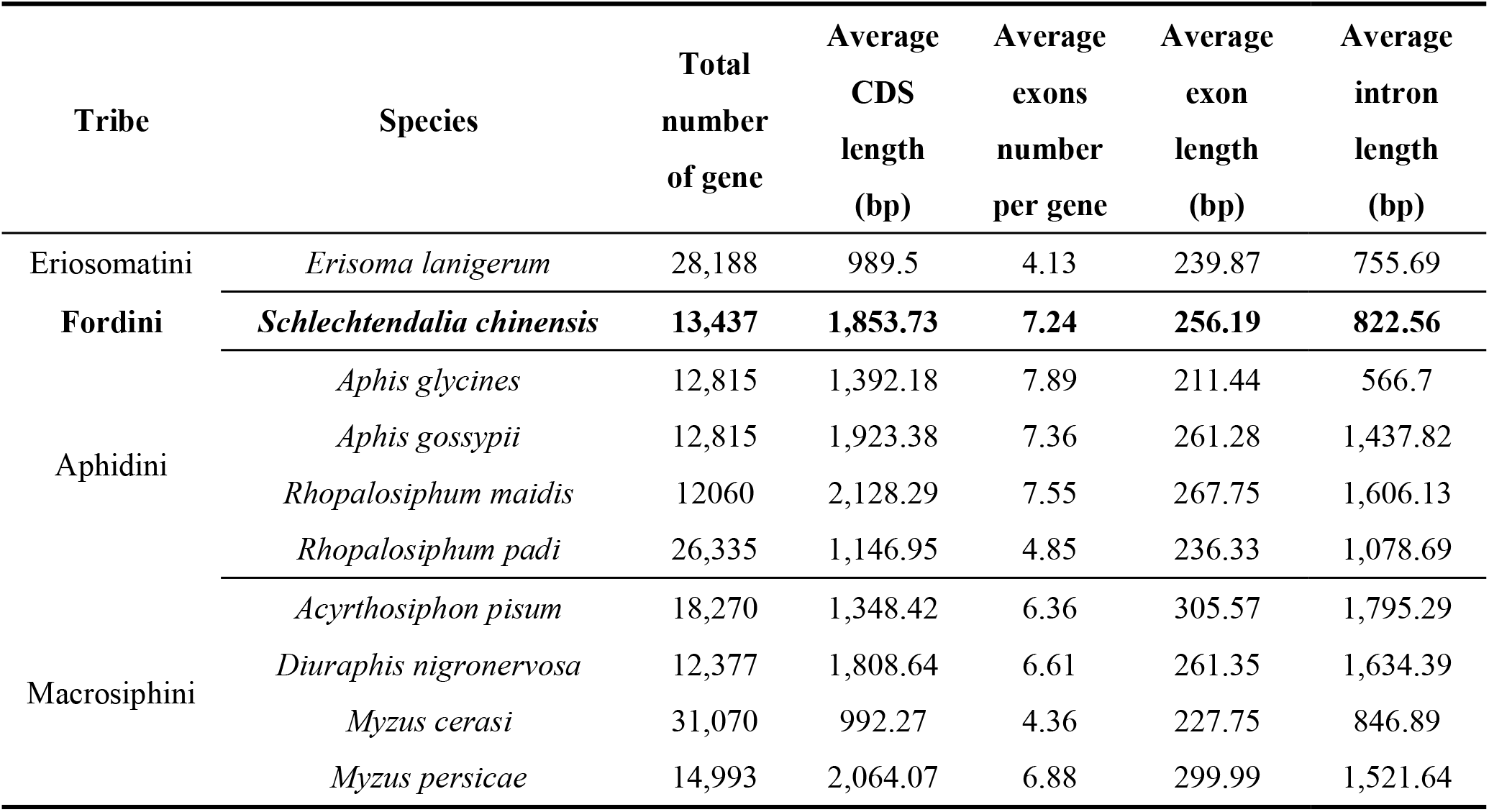
Summary statistics for predicted protein-coding genes and the number of P450 genes in *S. chinensis* and other nine aphids.

**Table 3.** Statistics for the functionally annotated protein-coding genes of *S. chinensis*.

| Type | database | Number | Percent (%) |
| --- | --- | --- | --- |
| Annotation | Nr | 11,457 | 85.26% |
|  | Nt | 7,687 | 57.21% |
|  | SwissProt | 7,531 | 56.05% |
|  | KOG | 5,896 | 43.88% |
|  | eggNOG | 11,110 | 82.68% |
|  | CDD | 10,106 | 75.21% |
|  | Interpro | 10,175 | 75.72% |
|  | GO | 8,739 | 65.04% |
|  | KEGG | 6,866 | 51.10% |
| Total | Annotated | 12,026 | 89.50% |
|  | Gene | 13,437 |  |

Among the 13,437 predicted genes, we were able to functionally annotate 12,026 (89.50%) genes (Table 3). This was based on the combination of 8,739 (65.04%) genes found via GO database and 6,866 (51.10%) genes present in the KEGG database. Additionally, we identified non-coding RNAs in the *S. chinensis* genome, including 128 tRNAs, 32 rRNAs, 29 miRNAs, and 81 snRNAs (Table S6).

### 3.3 Phylogenomic analysis

To explore this new genome assembly in a phylogenetic context and to investigate gene family evolution across aphids, the proteome of *S. chinensis* (the complete set of annotated protein coding genes) was compared to the proteomes of 10 other aphid species with fully sequenced genomes. A total of 30,300 single copy homologous genes were identified using orthogonal method and used for further phylogenetic analysis. *S. chinensis* was closely related to the wooly apple aphid, *E. lanigerum*, however, this relationship was paraphyletic. Instead, *S. chinensis* appeared to be the sister taxon to the Aphidini (Figure 4), placed evolutionary between the Eriosomatini and the Aphidini. Cedar bark aphid *C. cedri*, which belong to Lachnini formed a single clade at the base position, suggesting that it is probably the ancestral taxa. The other eight species clustered into two clades in consistence with morphological classification, with four species formed an Aphidini clade, and the other four species formed a Macrosiphini clade (Figure 4). The divergence time of *S. chinensis* and the two Aphidini+Macrosiphini aphid clades was estimated to be 433.28 MYA (Figure 4).

**Figure 3.**
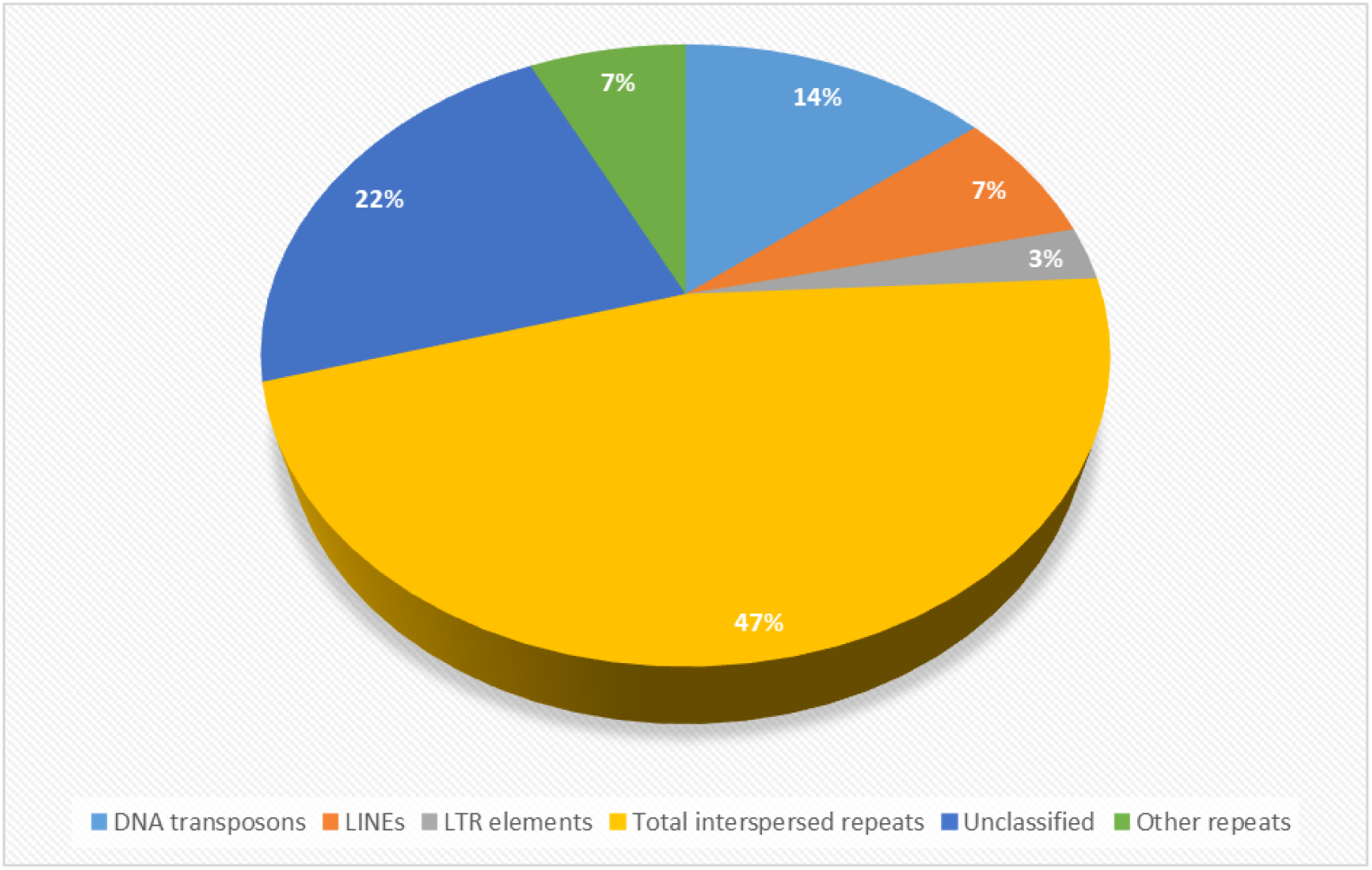
Repeats in the *S. chinensis* genome assembly.

**Figure 4.**
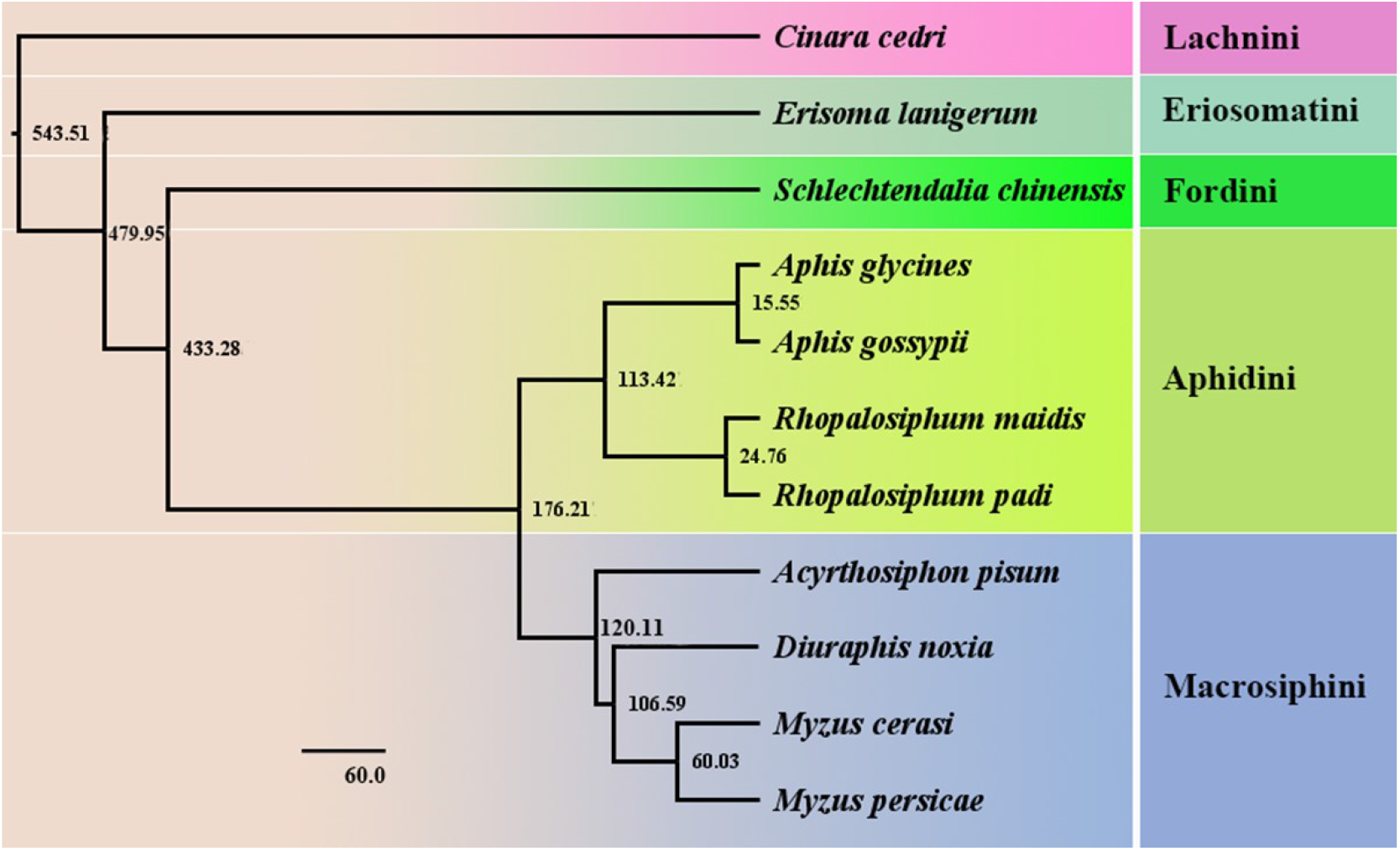
Timing of inferred divergence of *S. chinensis* and other ten aphids.

The results of gene-family expansion/constriction analysis showed a higher frequency of gene contraction in *S. chinensis*. Compared with the other ten aphid species, 228 gene families in *S. chinensis* underwent expansion, while 2451 gene families in *S. chinensis* underwent contraction (Figure S3). The roles of the expanded gene families appear to cell growth and cell death, lipid metabolism, the endocrine, nervous, immune system and signal transduction, using KEGG term enrichment analysis (Figure S4). While using GO pathway enrichment analysis, the roles of the expansion genes were related to response to stimulus, biological regulation, locomotion, catalytic activity and binding (Figure S5).

### 3.4 Salivary protein-encoding genes and gall formation associated genes

Plant galls are outgrowth induced by galling microorganisms and arthropods. Galling insects include galling aphids, midges and wasps. Previous studies have shown that gall induction is highly species-specific and galling insects deliver effectors into plant tissues, resulting in gall formation. However, the underlying mechanisms remained unknown. The gall midge *Mayetiola destructor* can inject effector proteins into tissues through its saliva during feeding, and convert a whole wheat seedling into a gall (Wang et al., 2018; Aljbory et al., 2020). *S. chinensis* feeds on host leaves where it injects saliva into the host leaf cells, resulting in gall formation. A total of 141 proteins have been identified from its salivary glands by LC-MS/MS analysis (Yang et al., 2018). In comparison to salivary proteins from 10 other free-living Hemipterans, the presence of a high proportion of proteins with binding activity was noticeable, including DNA-, protein-, ATP-, and iron-binding proteins, which may all be involved in gall formation. From the RNA-Seq analysis, transcripts corresponding to 14 protein-encoding genes encoding previously identified salivary proteins shown high expression levels at the fundatrix stage (Table S7 and Figure S6). Six of the genes encode functionally known proteins, including glucose dehydrogenase, aminobutyrate transaminase, ion transfer receptor, asparaginase, DnaJ, heat shock protein, and hexokinase. These salivary proteins are potentially involved in gall formation and plant defense inhibition (Table S7).

### 3.5 Cytochrome P450 genes

Cytochrome P450s play a central role in adaptation to plant chemicals and metabolizing plant metabolites (Li et al., 2019). Genome-wide analysis of P450 gene clusters providing an information on metabolic roles of orphan P450s (Xu, Wang & Guo, 2015). In the *S. chinensis* genome, we identified 36 P450 genes representing 0.27% of total genes. In comparison to the other nine aphid species, the genome of *S. chinensis* has the fewest P450 genes (Table S4). *S. chinensis* is an oligophagous insect, feeding only on the shrub *R. chinensis* in summer and on a few moss species in winter. By comparison, the other aphid species with sequenced genomes such as *M. persicae* are polyphagous, feeding on hundreds of host plants belonging to dozens families. We hypothesize that polyphagous aphids require more P450 genes (which range from 53 to 109) to detoxify the higher range of toxic compounds present in highly divergent host ranges (Table S4). Similar results from other insects also indicated that the oligophagous insect *Bombyx mori* had 70 P450 genes which less than the polyphagous insects, such as *Drosophila melanogaster* with 91 P450 genes, *Anopheles gambiae* with 112 P450 genes, and *Tribolium aegypti* with 164 P450 genes (d’Alencon et al., 2010; Ai et al., 2011). Polyphagous insects appeared to maintain much more cytochrome P450 gene numbers, likely due to the need for prompt response to the relative complex environmental challenges (Feyereisen, 2006).

In order to explore the roles of in *S. chinensis* for adaption to summer and winter host plants, gene expression profiles between the fundatrix and overwintering nymphs were analyzed using RNA-Seq. Our results showed that P450 transcript profiles were significantly different between aphid individuals feeding on the *R. chinensis* and mosses, suggesting that these genes are likely involved in host adaptation. All P450 genes were clustered into two significant groups. The first clade includes family members from chr01.3197 to chr10.020, which were more highly expressed in fundatrix than in overwintering nymphs. In contrast, the second clade includes family members from chr09.301to chr14.137, which were more highly expressed in overwintering nymphs than in fundatrix (Figure 5).

**Figure 5.**
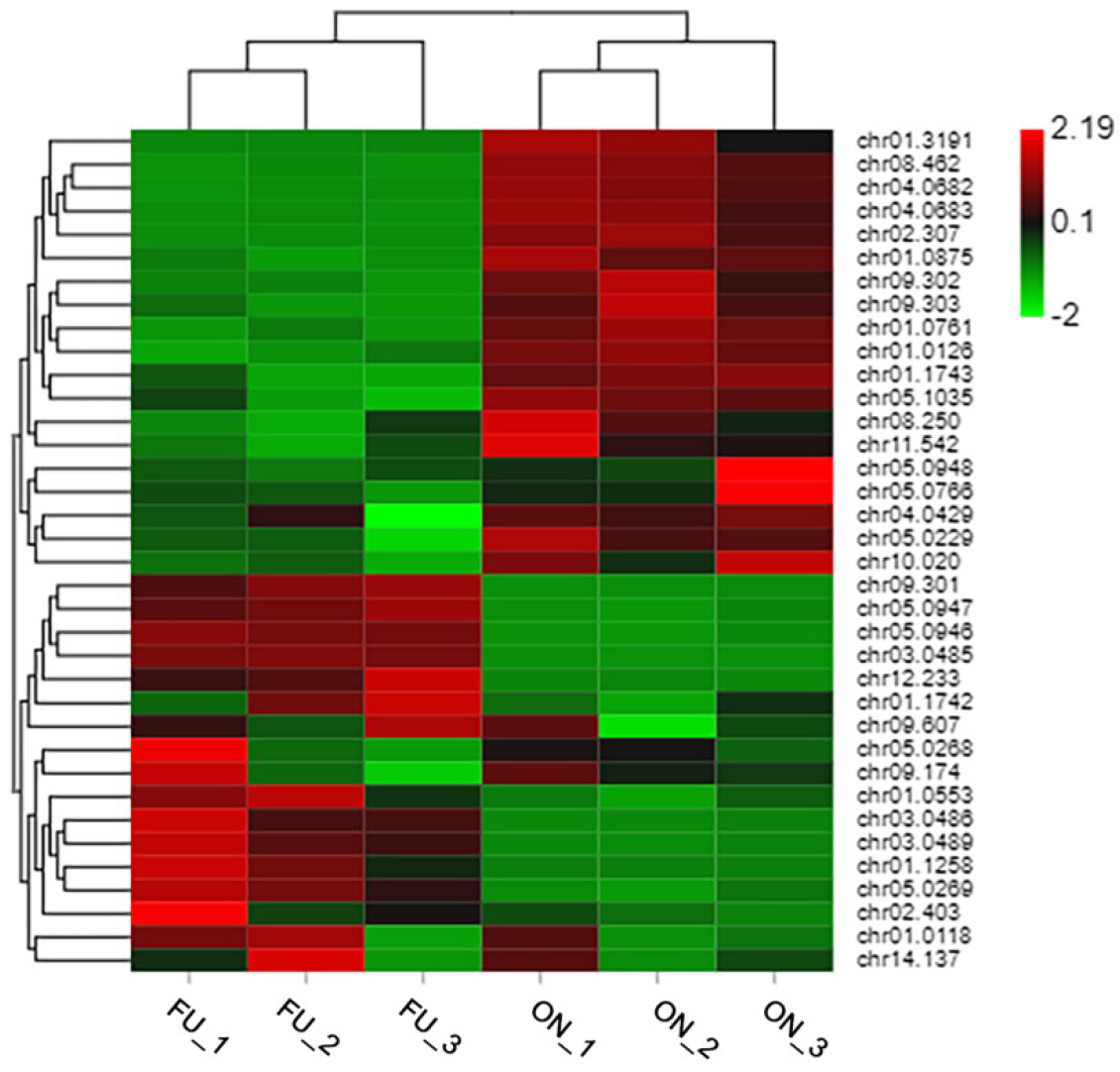
Expression of P450 in fundatrix and overwintering nymphs of *S. chinensis*. Abbreviation: FU: Fundatrix; ON: overwintering nymphs.

## Conclusion

We provided a high-quality chromosome-level genome assembly of the galling aphid *S. chinensis*. This represents the first genome sequence of an aphid that provides human benefits instead of an agriculture pest. The availability of the genome sequence should benefit future research on elucidating the molecular basis for the unique biology associated with the galling aphid and on revealing the molecular mechanisms for gall induction and host interactions. The availability of the genome sequence also provides the basis for potential genetic manipulation of the aphid for greater economic benefits to humans.

## Acknowledgements

We thank to Prof. Kirst King-Jones, University of Alberta, for his helpful comments.

## Author contributions

Z.C.X., Y.Z.X. and C.X.M conceived and designed the study. Z.C.X., Y.Z.X and W.H.Y. collected samples. W.H.Y. and Y.Y.X. performed the genome assembly, gene model prediction, gene annotation and comparative analyses. H.H.J. performed the chromosome analyses. W.H.Y., Y.Z.X. performed the transcriptome analyses. W.H.Y., Y.Z.X. and Y.Y.X. wrote the manuscript with input from all authors. Z.C.X., Y.Z.X. and C.M.S. analyzed the data and discussed the results. All authors reviewed the manuscript.

## Data availability statement

All data mentioned in this paper have been deposited in the National Center for Biotechnology Information with the BioProject accession number PRJNA700780 (genomic sequencing) and PRJNA702264 (transcriptome sequencing).

## Supplemental Information

**Table S1.** Sequencing data used for *S. chinensis* genome assembly.

|  | Illumina read | PacBio read | Hi-C |
| --- | --- | --- | --- |
| Raw data (Gb) | 73.85 | 130 | 100.09 |
| Clean data (Gb) | 71.94 | 121 | 96.23 |

**Table S2.** Statistical information of 19-kmer analysis.

| K | Heterozygosity | Genome<br>Haploid<br>Length (bp) | Genome<br>Repeat<br>Length (bp) | Genome<br>Unique<br>Length (bp) | Model<br>Fit | Read<br>Error<br>Rate |
| --- | --- | --- | --- | --- | --- | --- |
| 19 | 0.89% | 280,379,219 | 75,056,158.5 | 205,323,060.5 | 85.87% | 0.54% |

**Table S3.** Genome completeness assessment by BUSCO.

| BUSCO benchmark | Percentage (%) |  |  |  |
| --- | --- | --- | --- | --- |
|  | Eukaryota | Arthropoda | Insecta | Hemiptera |
| **Total BUSCO groups searched** | 255 | 1,013 | 1,367 | 2,510 |
| **Complete BUSCOs** | 91.80% | 96.40% | 94.60% | 97.80% |
| **Complete and single-copy BUSCOs** | 90.60% | 95.20% | 93.70% | 96.70% |
| **Complete and duplicated BUSCOs** | 1.20% | 1.20% | 0.90% | 1.10% |
| **Fragmented BUSCOs** | 3.90% | 1.20% | 1.50% | 0.60% |
| **Missing BUSCOs** | 4.30% | 2.40% | 3.90% | 1.60% |

**Table S4.** The comparisons of estimated genomes and P450 genes between *S. chinensis* and other nine aphids.

| Subfamily | Tribe | Species | Genome size (Mb) | Scaffold N50 | P450 genes | P450 gene Percentage (%) |
| --- | --- | --- | --- | --- | --- | --- |
| Eriosomatinae | Eriosomatini | *Eriosoma lanigerum* | 327 | 71.00 | 53 | 0.19 |
|  | Fordini | *Schlechtendalia chinensis* | 280 | 20.58 | 36 | 0.27 |
| Aphidinae | Aphidini | *Aphis glycines* | 303 | 0.174 | 58 | 0.30 |
|  |  | *Aphis gossypii* | 294 | 0.438 | 73 | 0.57 |
|  |  | *Rhopalosiphum maidis* | 326 | 0.900 | 75 | 0.62 |
|  |  | *Rhopalosiphum padi* | 319 | 0.116 | 58 | 0.48 |
|  | Macrosiphini | *Acyrtosiphon pisum* | 525 | 126.6 | 109 | 0.60 |
|  |  | *Diuraphis noxia* | 393 | 0.397 | 66 | 0.50 |
|  |  | *Myzus cerasi* | 406 | 0.023 | 67 | 0.23 |
|  |  | *Myzus persicae* | 295 | 69.48 | 80 | 0.34 |

**Table S5.** Summary of chromosome in the *S. chinensis*.

| Chromosome | Length (bp) |
| --- | --- |
| chr01 | 86,700,217 |
| chr02 | 21,093,922 |
| chr03 | 20,851,048 |
| chr04 | 20,581,381 |
| chr05 | 19,598,981 |
| chr06 | 19,543,707 |
| chr07 | 14,915,226 |
| chr08 | 13,343,717 |
| chr09 | 11,956,905 |
| chr10 | 11,438,933 |
| chr11 | 11,114,567 |
| chr12 | 10,874,994 |
| chr13 | 10,139,000 |
| Total anchored | 272,152,598 |
| Unanchored | 8,300,227 |
| Percentage | 97% |

**Table S6.**
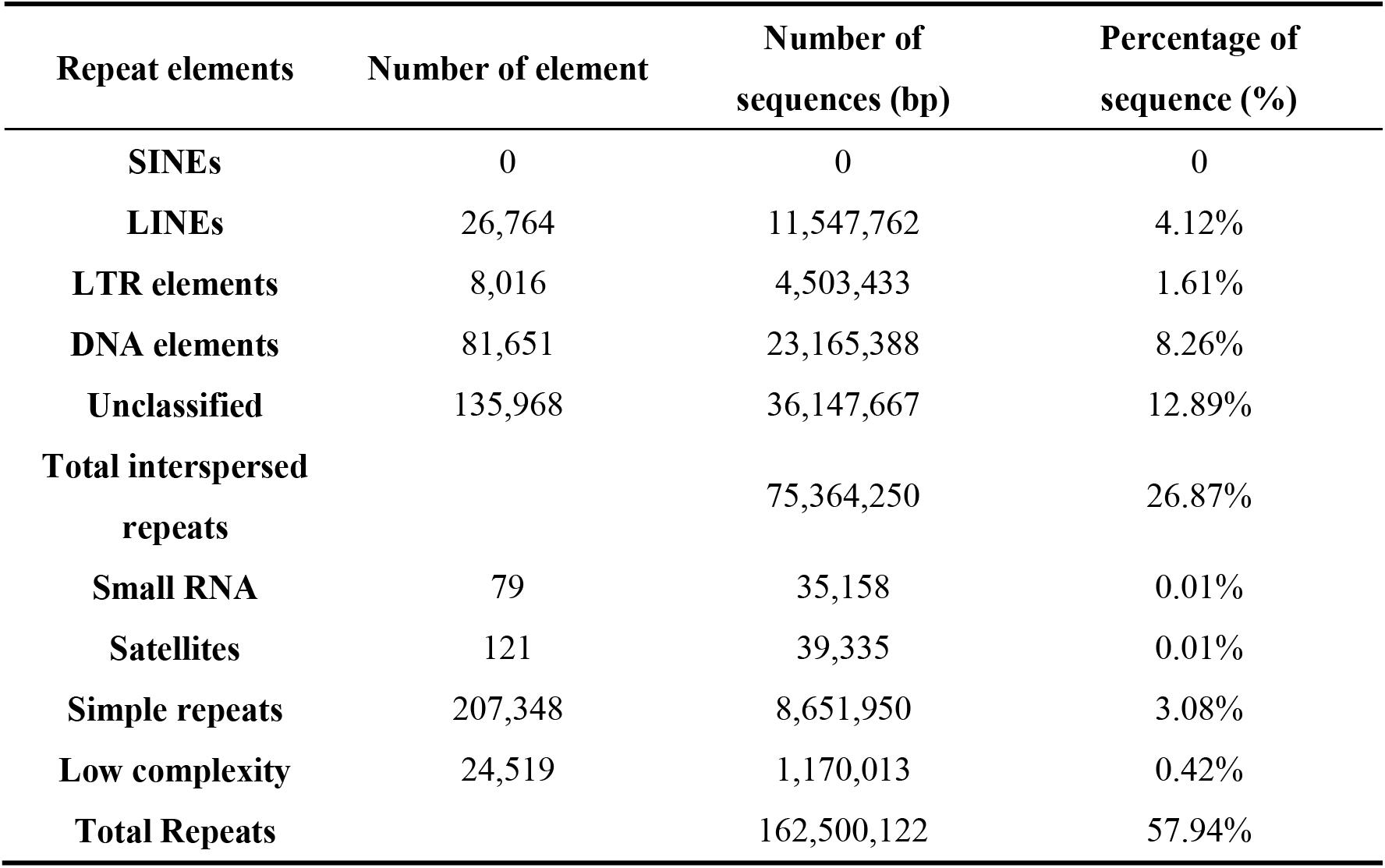
Repeats in the *S. chinensis* genome assembly.

| Repeat elements | Number of element | Number of sequences (bp) | Percentage of sequence (%) |
| --- | --- | --- | --- |
| SINEs | 0 | 0 | 0 |
| LINEs | 26,764 | 11,547,762 | 4.12% |
| LTR elements | 8,016 | 4,503,433 | 1.61% |
| DNA elements | 81,651 | 23,165,388 | 8.26% |
| Unclassified | 135,968 | 36,147,667 | 12.89% |
| Total interspersed repeats |  | 75,364,250 | 26.87% |
| Small RNA | 79 | 35,158 | 0.01% |
| Satellites | 121 | 39,335 | 0.01% |
| Simple repeats | 207,348 | 8,651,950 | 3.08% |
| Low complexity | 24,519 | 1,170,013 | 0.42% |
| Total Repeats |  | 162,500,122 | 57.94% |

**Table S7.**
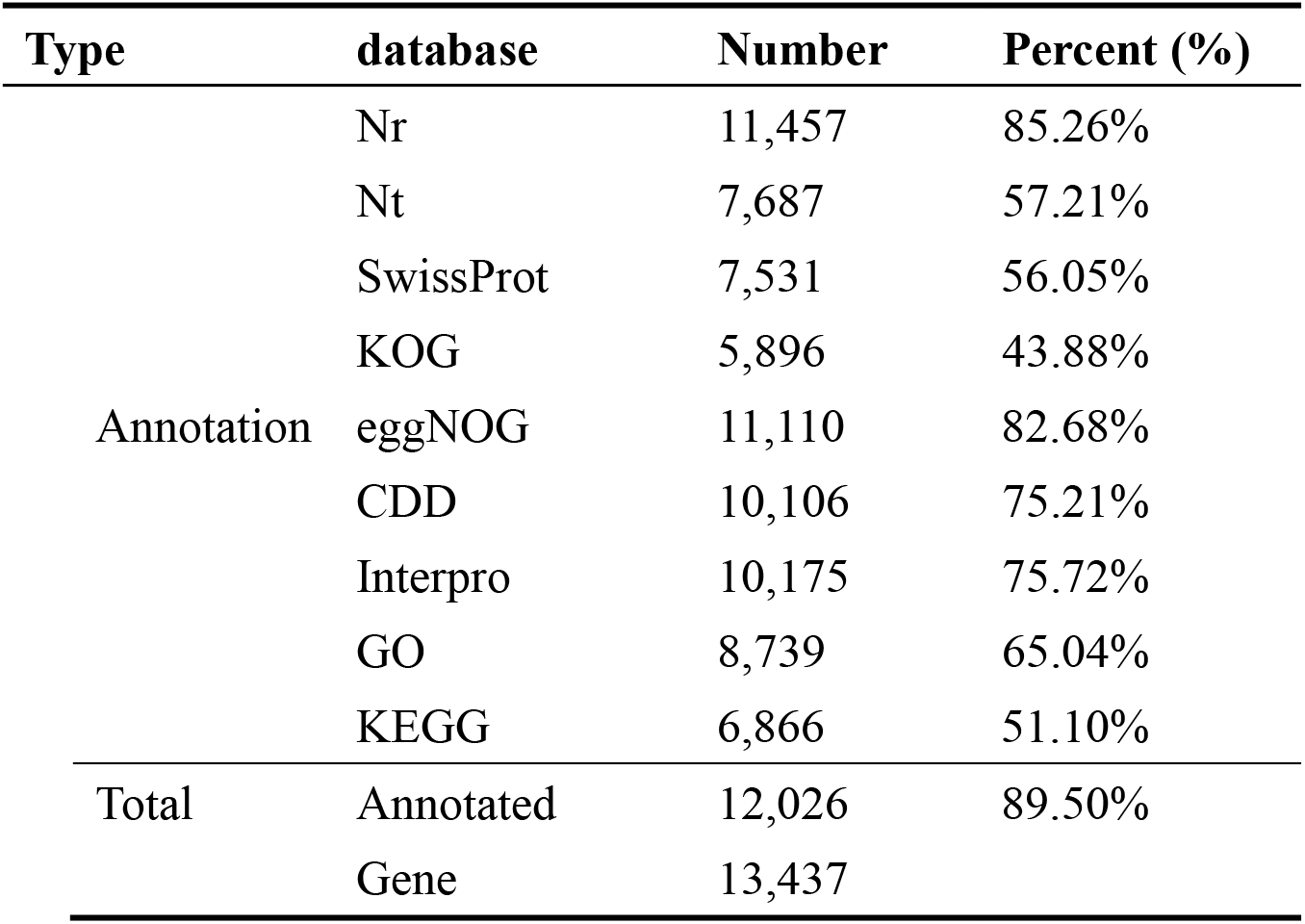
Statistics for the functionally annotated protein-coding genes.

**Table S8.** Functional annotation of salivary proteins specifically expressed in fundatrix.

| Gene ID | Name of coding protein |
| --- | --- |
| **chr01.2056** | Ankyrin |
| **chr13.018** | Glucose dehydrogenase |
| **chr01.0223** | Glucose dehydrogenase |
| **chr01.1395** | Uncharacterized protein |
| **chr01.2438** | Uncharacterized protein |
| **chr03.0480** | Aminobutyrate transaminase |
| **chr05.0056** | Actin |
| **chr02.167** | Ion transfer receptor |
| **chr08.259** | Asparaginase |
| **chr03.0139** | Tubulin |
| **chr06.0990** | Calphotin |
| **chr09.570** | DnaJ homologous subfamily |
| **chr03.0155** | Heat shock protein |
| **chr02.176** | Hexokinase |

**Figure S1.**
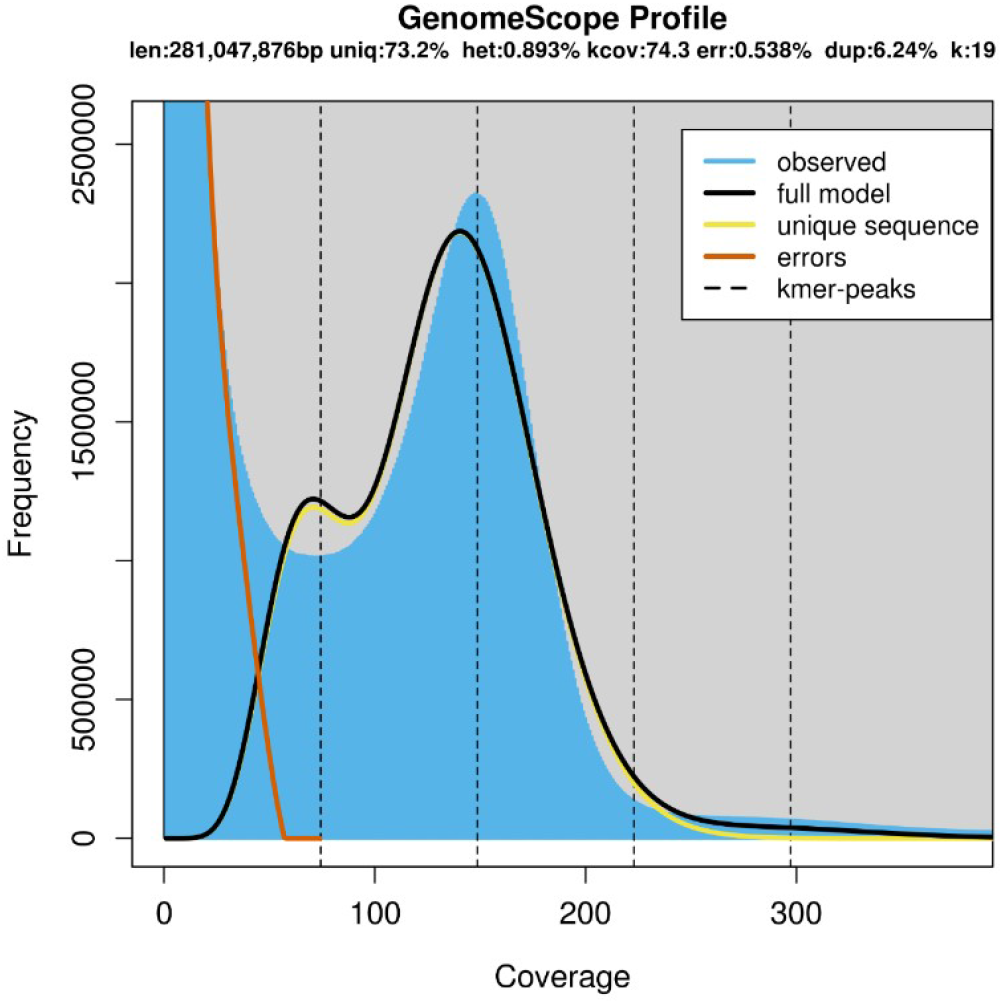
Distribution frequency of 19-mers in the *S. chinensis* genome.

**Figure S2.**
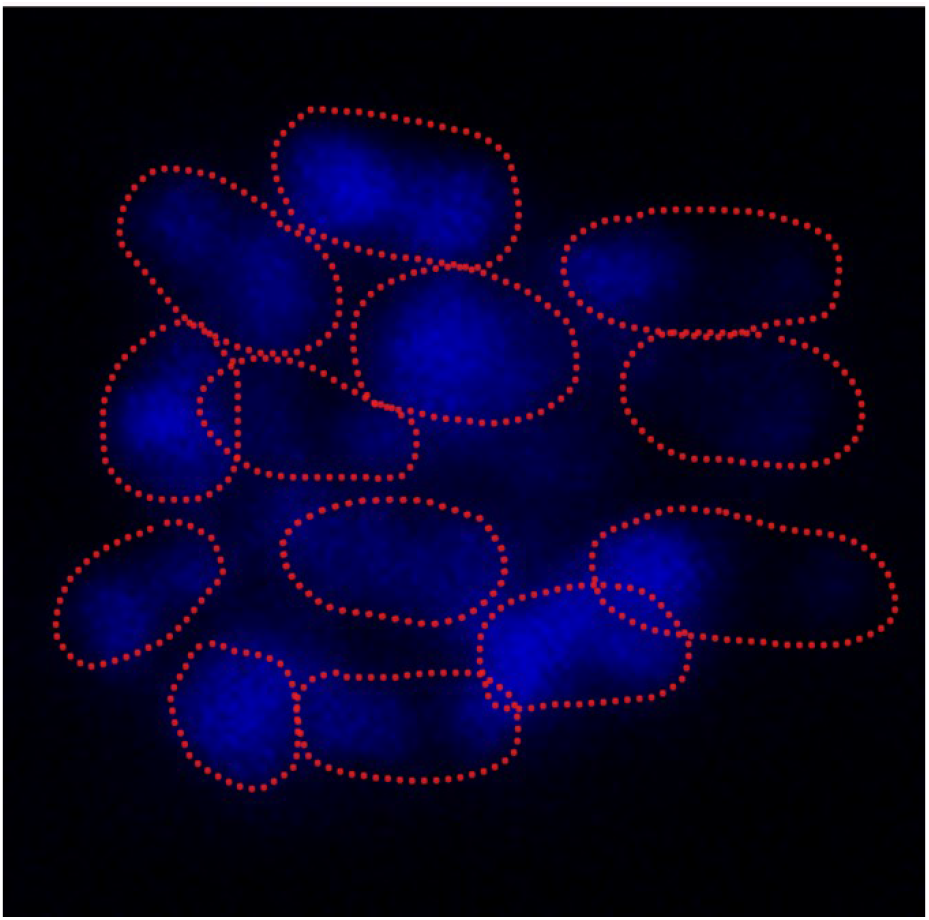
Chromosomic staining of fundatrigenia salivary gland cells. The red signal indicates the condensed chromosomes.

**Figure S3.**
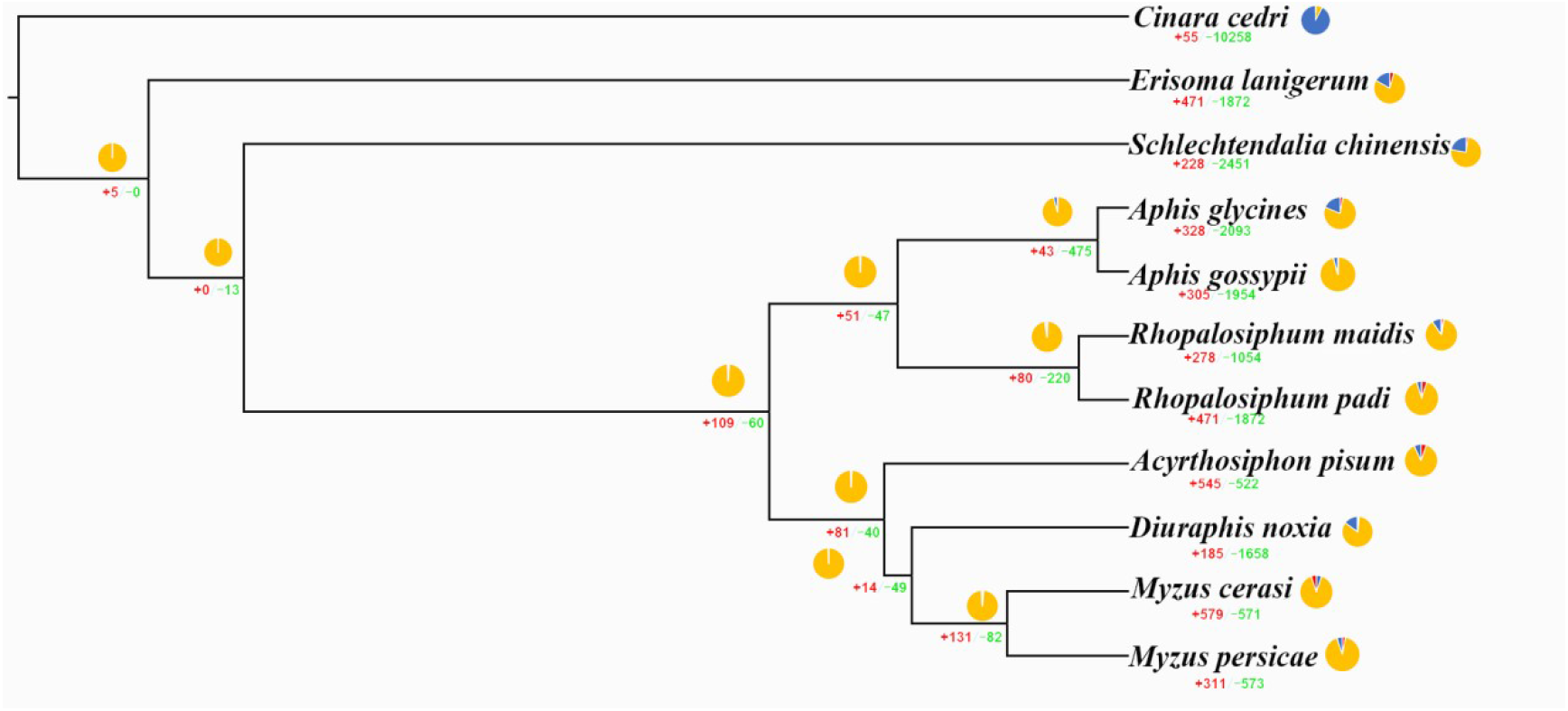
Gene family evolutions in *S. chinensis* and other ten aphids. The pie charts represent proportions of gene family expansions, contractions, or no changes. Gene family expansions are indicated inred. Gene family contractions are indicated in blue. Gene families with no changes are indicated in yellow.

**Figure S4.**
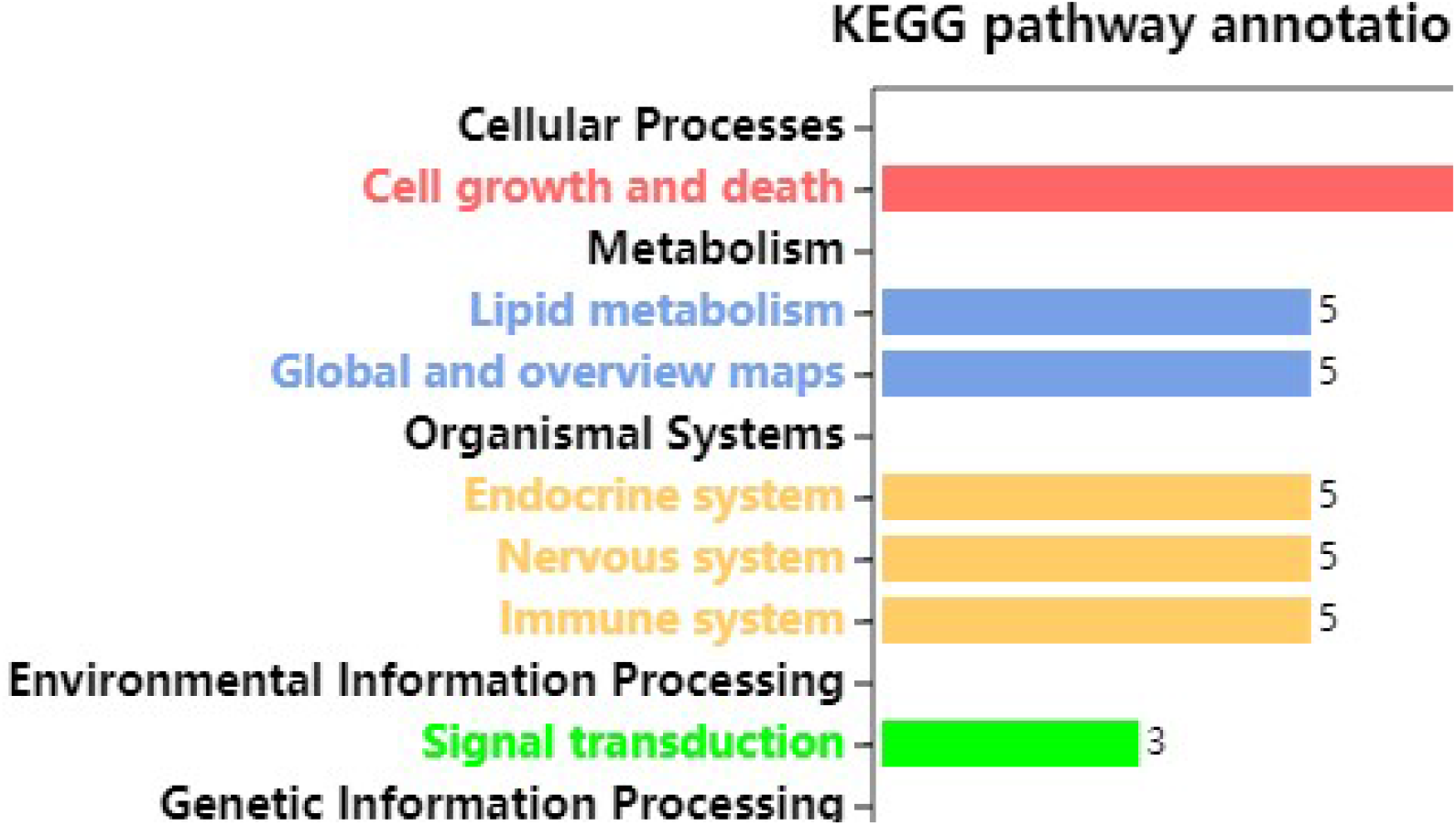
KEGG pathway enrichment analysis was performed for the expansion gene family of *S. chinensis*. The graph depicts the most highly enriched pathways.

**Figure S5.**
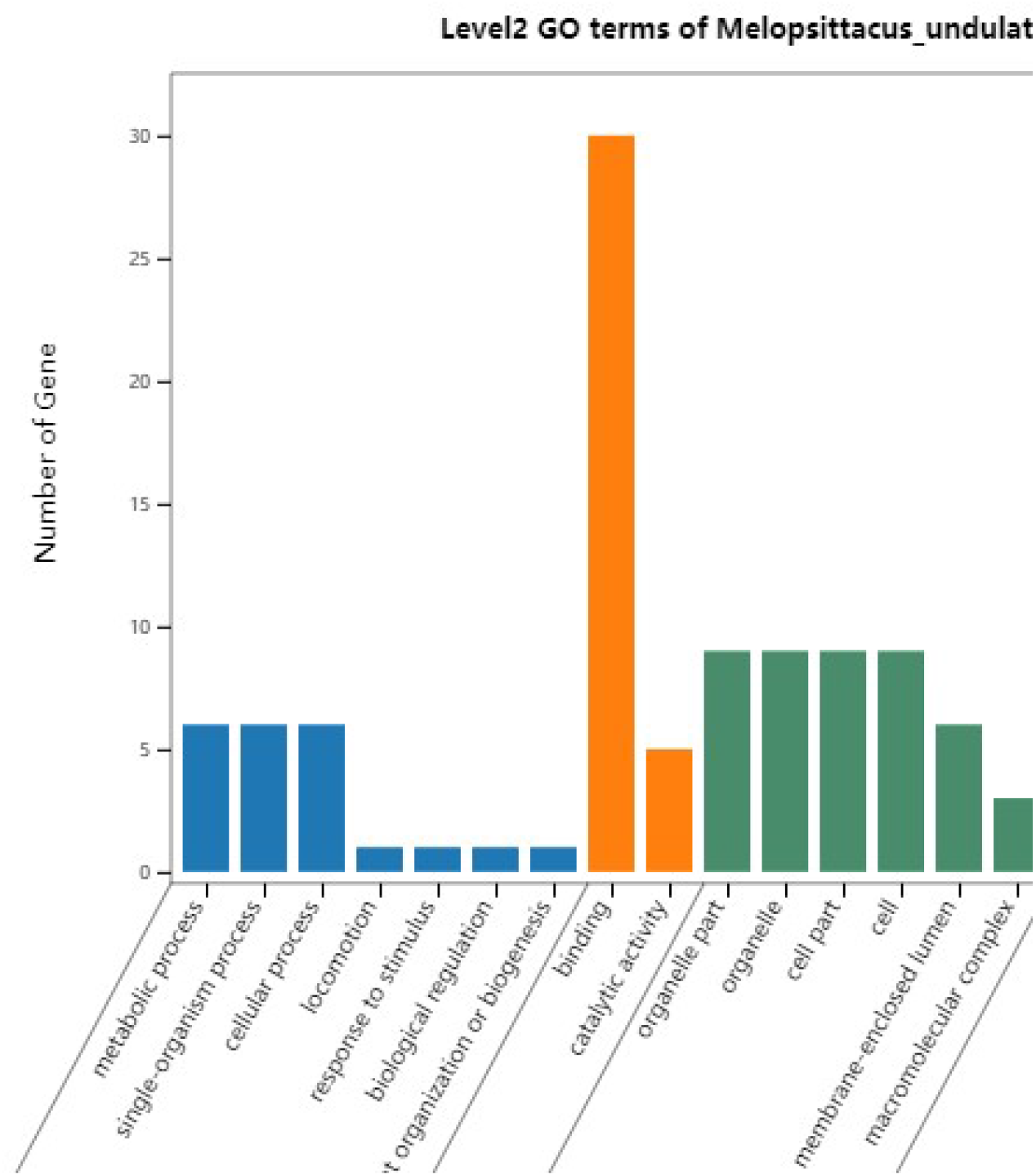
Gene ontology (GO) enrichment analysis of expanded gene families of *S. chinensis*. Bars are subdivided to representdifferent GO terms. The top 20 most significant GO categories were included (p <0.05). BP, biological process; CC, cellular component; MF, molecular function.

**Figure S6.**
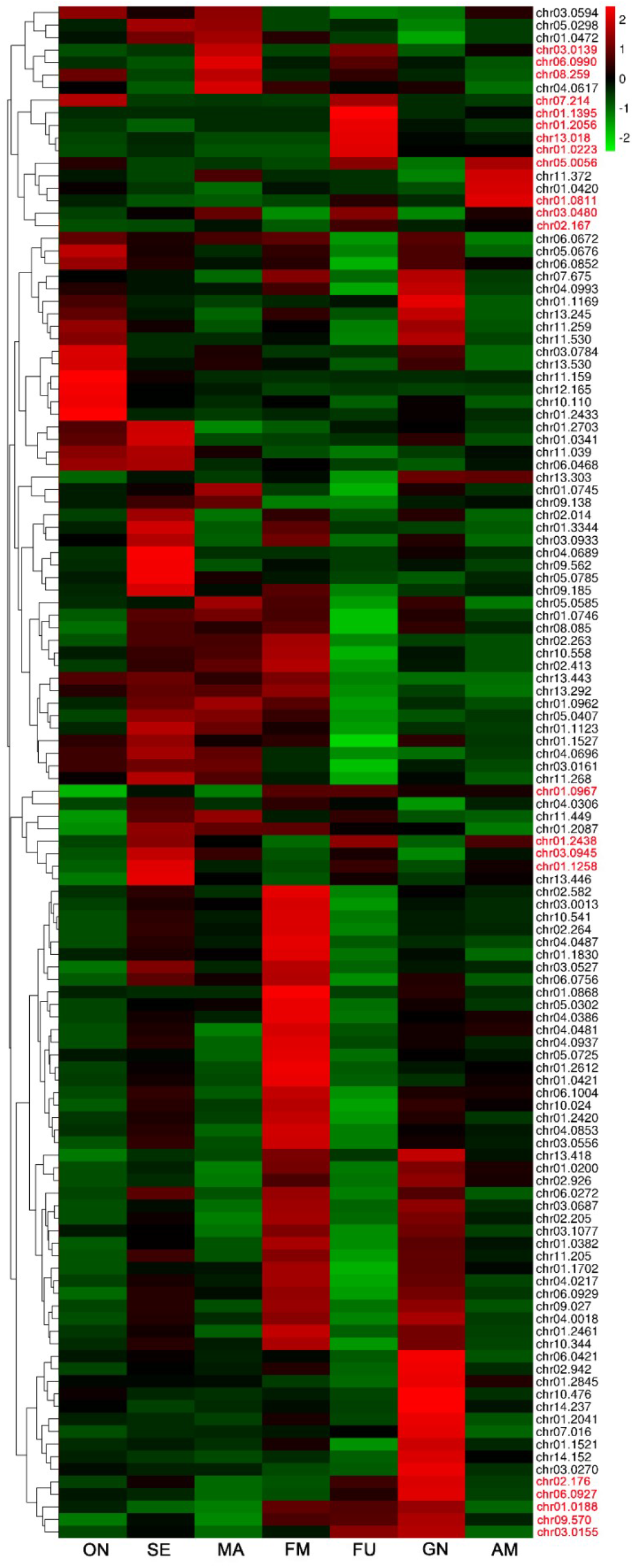
The expression of salivary proteins in each stage of *S. chinensis*. Abbreviation: ON: Overwintering nymphs; SE: Sexuparae (spring migrant); MA: Male; FM: Female; FU: Fundatrix; GN: Gall nymphs (fundatrigeniae); AM: Autumn migrant.

## References

Ai, J. W., Zhu, Y., Duan, J., Yu, Q. Y., Zhang, G. J., Wan, F., & Xiang, Z. H. (2011). Genome-wide analysis of cytochrome P450 monooxygenase genes in the silkworm, Bombyx mori. Gene, 480, 42–50. https://doi.org/10.1016/j.gene.2011.03.002

Alibory, Z., Micheal, J. A., Park, Y., Reeck, G. R., & Chen M. S. (2020). Differential localization of Hessian fly candidate effectors in resistant and susceptible wheat plants. Plant direct, 00: 1–15. https://doi.org/10.1002/pld3.246

Biello, R., Singh, A., Godfrey, C. J., Fernández, F. F., Mugford, S. T., & Powell, & G., Hogenhout, S. A., | Mathers, C. (2021). A chromosome level genome assembly of the woolly apple aphid, Eriosoma lanigerum Hausmann (Hemiptera: Aphididae). Molecular Ecology Resources, 21(1), 316–326. https://doi.org/10.1111/1755-0998.13258

Birney, Ewan, Clamp, Michele, Durbin, & Richard. (2004). GeneWise and Genomewise. Genome Research, 14(5), 988–995. https://doi.org/10.1101/gr.1865504

Blackman, R. L., & Eastop, V. F. (2017). Taxonomic issues. In H. F. van Emden, & R. Harrington (Eds.), Aphids as crop pests (2nd ed.). CAB International.

Blackman, R. L., & Eastop, V. F. (2000). Aphids on the world’s crops: An identification and information guide. John Wiley & Sons Ltd.

Blanco, E., Parra, G., & Guigó, R. (2002). Using geneid to Identify Genes. John Wiley & Sons, Inc.

Castresana, J. (2000). Selection of conserved blocks from multiple alignments for their use in phylogenetic analysis. Molecular Biology Evolution, 17(4), 540–552. https://doi.org/10.1093/oxfordjournals.molbev.a026334

Chen, X. M., Yang, Z. X., Chen, H., Qi, Q., Liu, J., & Wang, C., Shao, S. X., Lu, Q., Li, Y., Wu, H. X., King-Jones, K., Chen, M. S. (2020). A complex nutrient exchange between a gall-forming aphid and its plant host. Frontiers in Plant Science, 11, 811. https://doi.org/10.3389/fpls.2020.00811

D’Alençon, E., Sezutsu, H., Legeai, F., Permal, E., Bernard-Samain, S., Gimenez, S…. & Feyereisen, R. (2010). Extensive synteny conservation of holocentric chromosomes in Lepidoptera despite high rates of local genome rearrangements. PNAS, 107(17), 7680–7685. https://doi.org/10.1073/pnas.0910413107

Dudchenko, O., Batra, S. S., Omer, A. D., Nyquist, S. K., Hoeger, M., Durand, N. C., Shamim, M. S., Machol, I.P., Lander, E. S., Aiden, A. P., & Aiden, E. L. (2017). De novo assembly of the Aedes aegypti genome using Hi-C yields chromosome-length scaffolds. Science, 356, 92–95. https://doi.org/10.1126/science.aal3327

Feyereisen, R. (2006). Evolution of insect P450. Biochemistry Society Transition, 34(6), 1252–1255. https://doi.org/10.1042/BST0341252

Flynn, J. M., Hubley, R., Goubert, C., Rosen, J., Clark, A. G., Feschotte, C., & Smit, A. F. (2020). RepeatModeler2 for automated genomic discovery of transposable element families. PNAS, 117(17), 9451–9457. https://doi.org/10.1073/pnas.1921046117

Haas, B. J., Salzberg, S. L., Zhu, W., Pertea, M., Allen, J. E., & Orvis, J., White, O., Buell, C. R., & Wortman J. R. (2008). Automated eukaryotic gene structure annotation using EVidenceModeler and the Program to Assemble Spliced Alignments. Genome biology, 9(1), R7. https://doi.org/10.1186/gb-2008-9-1-r7

Hahn, M. W., Demuth, J. P., & Han, S. G. (2007). Accelerated rate of gene gain and loss in primates. Genetics, 177(3). https://doi.org/10.1534/genetics.107.080077

Hirano, T., Kimura, S., Sakamoto, T., Okamoto, A., Nakayama, T., Matsuura, T., Ikeda, Y., Takeda, S., Suzuki, Y., Ohshima I., & Sato, M. H. Reprogramming of the developmental program of Rhus javanica during initial stage of gall induction by Schlechtendalia chinensis. Frontiers in Plant Science, 2020, 11, 471. https://doi.org/10.3389/fpls.2020.00471

International Aphid Genomics Consortium. (2010). Genome Sequence of the Pea Aphid Acyrthosiphon pisum. Plos Biology, 8(2), 1–25. https://doi.org/10.1371/journal.pbio.1000313

Julca, I., Marcet-Houben, M., Cruz, F., Vargas-Chavez, C., Johnston, J. S., Gómez-Garrido, J., Frias, L., Corvelo, A., Loska, D., Cámara, F., Gut, M., Alioto, T., Latorre, A., & Gabaldón, T. (2020). Phylogenomics identifies an ancestral burst of gene duplications predating the diversification of aphidomorpha. Molecular Biology and Evolution, 37(3), 730–756. https://doi.org/10.1093/molbe v/msz261

Hu, J., Fang, J. P., Su, Z. Y., & Liu, S. L. (2019). NextPolish: a fast and efficient genome polishing tool for long-read assembly. Bioinformatics, (7), 7. https://doi.org/10.1093/bioinformatics/btz891

Huang, S. F., Kang, M. J., & Xu, A. L. (2017). HaploMerger2: rebuilding both haploid sub-assemblies from high-heterozygosity diploid genome assembly. Bioinformatics 16, 2577. https://doi.org/10.1093/bioinformatics/btx220

Katoh, K., Misawa, K., Kuma, K., & Miyata, T. (2002). MAFFT: a novel method for rapid multiple sequence alignment based on fast Fourier transform. Nucleic Acids Research, 30(14), 3059–3066. https://doi.org/10.1093/nar/gkf436

Katoh, K., & Standley, D. (2013). MAFFT multiple sequence alignment software version 7: improvements in performance and usability. Molecular Biology and Evolution, 30(4), 772–780. https://doi.org/10.1093/molbev/mst010

Kalvari, I., Argasinska, J., Quinones-Olvera, N., Nawrocki, E. P., Rivas, E., Eddy, S. R., Bateman, A., Finn, R. D., & Petrov, A. I. (2018). Rfam 13.0: shifting to a genome-centric resource for non-coding RNA families. Nucleic Acids Research 46 (Database issue), D335–D342. http://dx.doi.org/10.1093/nar/gkx1038

Li, F., Zhao, X., Li, M., He, K., Huang, C., Zhou, Y., Li, Z., & Walters, J. R. (2019). Insect genomes: progress and challenges. Insect Molecular Biology, 28(6), 739–758. https://doi.org/10.1111/imb.12599

Li, L., Stoeckert, C. J., & Roos, D. (2003). OrthoMCL: identification of ortholog groups for eukaryotic genomes. Genome Research, 13(9), 2178–2189. https://doi.org/10.1101/gr.1224503

Liu, P., Yang, Z. X., Chen, X. M., Foottit, R. G. (2014). The Effect of the gall-forming aphid Schlechtendalia chinensis (Hemiptera: Aphididae) on leaf wing ontogenesis in Rhus chinensis (Sapindales: Anacardiaceae). Annals of the Entomological Society of America, 107(1), 242–250. http://www.bioone.org/doi/full/10.1603/AN13118

Li, Y., Park, H., Smith, T. E., & Moran, N. A. (2019). Gene family evolution in the pea aphid based on chromosome-level genome assembly. Molecular Biology and Evolution, 36(10), 2143–2156. https://doi.org/10.1093/molbev/msz138

Mathers, T. C. (2020). Improved genome assembly and annotation of the soybean aphid (Aphis glycines Matsumura). G3: Genes, Genomes, Genetics, 10(3), g3.400954.2019. https://doi.org/10.1534/g3.119.400954

Mathers, T. C., Chen, Y., Kaithakottil, G., Legeai, F., Mugford, S. T., Baa-Puyoulet, P., Bretaudeau, A., Clavijo, B., Colella, S., Collin, O., Dalmay, T., Derrien, T., Feng, H., Gabaldón, T., Jordan, A., Julca, I., Kettles, G. J., Kowitwanich, K., Lavenier, D., … Hogenhout, S. A. (2017). Rapid transcriptional plasticity of duplicated gene clusters enables a clonally reproducing aphid to colonise diverse plant species. Genome Biology, 18(1), 27. https://doi.org/10.1186/s13059-016-1145-3

Mathers, T. C., Mugford, S. T., Hogenhout, S. A. T., & Tripathi, L. (2020). Genome sequence of the banana aphid, Pentalonia nigronervosa Coquerel (Hemiptera: Aphididae) and its symbionts. G3: Genes, Genomes, Genetics, 10(12), 4315–4321. https://doi.org/10.1534/g3.120.401358

Mathers, T. C., Wouters, R. H. M., Mugford, S. T., Swarbreck, D., Van Oosterhout, C., & Hogenhout, S. A. (2020). Chromosome-scale genome assemblies of aphids reveal extensively rearranged autosomes and long-term conservation of the X chromosome. Molecular Biology and Evolution, https://doi.org/10.1093/molbev/msaa246

Moran, N. A. (1989). A 48-million-year-old aphid-host plant association and complex life cycle: biogeographic evidence. Science, 245(4914), 173–175. https://doi.org/10.1126/science.245.4914.173

Nicholson, S. J., Nickerson, M. L., Dean, M., Song, Y., Hoyt, P. R., Rhee, H., Kim, C., & Puterka, G. J. (2015). The genome of Diuraphis noxia, a global aphid pest of small grains. BMC Genomics, 16(1), 1–16. https://doi.org/10.1186/s12864-015-1525-1

Quan, Q. M., Hu, X., Pan, B. H., Zeng, B. S., Wu, N. N., Fang, G. Q., Cao, Y. H., Chen, X. Y., Li, X., Huang, Y. P., & Zhan, S. (2019). Draft genome of the cotton aphid Aphis gossypii. Insect Biochemistry and Molecular Biology, 105, 25–32. https://doi.org/10.1016/j.ibmb.2018.12.007

Rao, S. S. P., Huntley, M. H., Durand, N. C., Stamenova, E. K., Bochkov, I. D., Robinson, J. T., Sanborn, A. L., Machol, I., Omer, A. D., & Lander, E. S. (2014). A 3D map of the human genome at Kilobase resolution reveals principles of chromatin looping. Cell, 158, 1–6. https://doi.org/10.1016/j.cell.2014.11.021

Ranallo-Benavidez, T. R., Jaron, K. S., & Schatz, M. C. (2020). GenomeScope 2.0 and Smudgeplot for reference-free profiling of polyploid genomes. Nature Communications, 11(1), 1432. https://doi.org/10.1038/s41467-020-14998-3

Stamatakis, A. (2014). RAxML version 8: a tool for phylogenetic analysis and post-analysis of large phylogenies. Bioinformatics, 30(9), 1312–1313. https://doi.org/10.1093/bioinformatics/btu033

Stanke, M., Keller, O., Gunduz, I., Hayes, A., Waack, S., & Morgenstern, B. (2006). AUGUSTUS: ab initio prediction of alternative transcripts. Nucleic Acids Research, 34 (Web Server issue), W435–439. https://doi.org/10.1093/nar/gkl200

Thorpe, P., Escudero-Martinez, C. M., Cock, P. J. A., Eves-van den Akker, S., & Bos, J. I. B. (2018). Shared transcriptional control and disparate gain and loss of aphid parasitism genes. Genome Biology and Evolution, 10(10), 2716–2733. https://doi.org/10.1093/gbe/evy183

Takeda, S., Yoza, M., Amano, T., Ohshima, I., Hirano, T., Sato, M. H., Sakamoto, T., & Seisuke Kimura, S. (2019) Comparative transcriptome analysis of galls from four different host plants suggests the molecular mechanism of gall development. PLoS One, 14(10), e0223686. https://doi.org/10.1371/journal.pone.0223686

Walker, B. J., Abeel, T., Shea, T., Priest, M., Abouelliel, A., Sakthikumar, S., Cuomo, C. A., Zeng, Q. D., Wortman, J., Young, S. K., & Earl, A. M. (2014). Pilon: an integrated tool for comprehensive microbial variant detection and genome assembly improvement. PLoS One, 9(11), e112963. https://doi.org/10.1371/journal.pone.0112963

Wang, Z., Ge, J., Chen, H., Cheng, X., Yang, Y., Li, J., Whitworth, R. J., & Chen M. C. S. (2018). An insect nucleoside diphosphate kinase (NDK) functions as an effector protein in wheat - Hessian fly interactions. Insect Biochemistry and Molecular Biology, 100, 30–38. https://doi.org/10.1016/j.ibmb.2018.06.003

Wenger, J. A., Cassone, B. J., Legeai, F., Johnston, J. S., Bansal, R., Yates, A. D., Coates, B. S., Pavinato, V. A. C., & Michel, A. (2016). Whole genome sequence of the soybean aphid, Aphis glycines. Insect Biochemistry and Molecular Biology, 123, 102917. https://doi.org/10.1016/j.ibmb.2017.01.005

Wenger, J. A., Cassone, B. J., Legeai, F., Johnston, J. S., Bansal, R., Yates, A. D., Coates, B. S., Pavinato, V. A. C., & Michel, A. (2020). Whole genome sequence of the soybean aphid, Aphis glycines. Insect Biochemistry & Molecular Biology, 123, 102917. https://doi.org/10.1016/j.ibmb.2017.01.005

Wool D. Galling aphids: specialization, biological complexity, and variation. (2004). Annual Review of Entomology, 49(1), 175. https://doi.org/10.1146/annurev.ento.49.061802.123236

Xu, J., Wang, X., & Guo, W. (2015). The cytochrome P450 superfamily: Key players in plant development and defense. Journal of Integrative Agriculture, 14(9), 1673–1686. https://doi.org/10.1016/S2095-3119(14)60980-1

Yang, Z. H. (2007). PAML 4: phylogenetic analysis by maximum likelihood. Molecular Biology and Evolution, 24(8), 1586–1591. https://doi.org/10.1093/molbev/msm088

Yang Z. X., Ma, L., Francis, F., Yang, Y., Chen, H., Wu, H. X., & Chen, X. M. (2018). Proteins identified from saliva and salivary glands of the Chinese gall aphid Schlectendalia chinensis. Proteomics, 18, 1700378. https://doi.org/10.1002/pmic.201700378

Zhang, C. X., Tang, X. D., & Cheng, J. A. (2008). The utilization and industrialization of insect resources in China. Entomological research, 38, S38–S47. https://doi.org/10.1111/j.1748-5967.2008.00173.x

Zhang, G. X., Qiao, G. X., Zhong, T. S., & Zhang, W. Y. (1999). Fauna Sinica, Insecta Vol. 14 Homoptera, Mindaridae and Pemphigidae. Science Press, Beijing.

